# N-acetylcysteine and acetyl-L-carnitine did not prevent acute PTZ-induced seizure in adult and larvae zebrafish

**DOI:** 10.1101/2023.07.13.548882

**Authors:** Rafael Chitolina, Carlos G. Reis, Thailana Stahlhofer-Buss, Amanda Linazzi, Radharani Benvenutti, Matheus Marcon, Ana P. Herrmann, Angelo Piato

## Abstract

Epilepsy is a prevalent neurological disorder, affecting approximately 1 to 2% of the global population. The hallmark of epilepsy is the occurrence of epileptic seizures, which are characterized by predictable behavioral changes reflecting the underlying neural mechanisms of the disease. Unfortunately, around 30% of patients do not respond to the current available pharmacological treatments. Consequently, it is crucial to explore alternative therapeutic options for managing these seizures. Two potential candidates for attenuating seizures are N-acetylcysteine (NAC) and Acetyl-L-carnitine (ALC), as they have shown promising neuroprotective effects through the modulation of the neurotransmitter glutamate. Therefore, this study aims to assess the effects of varying concentrations (0.1, 1.0, and 10 mg/L) of NAC and ALC on acute PTZ-induced seizures in zebrafish, in both adult and larval stages. The evaluation of behavioral parameters such as seizure intensity and latency to the crisis can provide insights into the efficacy of these substances. However, our results indicate that both drugs at any of the tested concentrations were not able to reduce PTZ-induced epileptic seizures. On the other hand, the administration of diazepam demonstrated a notable reduction in seizure intensity and an increase in latencies to higher scores of epileptic seizures. Consequently, we conclude that, under the conditions employed in this study, NAC and ALC do not exhibit any significant effects on acute seizures in zebrafish.

## Introduction

Epilepsy is a prevalent neurological disorder that affects approximately 1 to 2% of the global population (Falco-Walter, 2020). It is characterized by the occurrence of two or more unprovoked or reflex seizures, with a minimum time interval of 24 hours between them. Alternatively, it can be defined as a single unprovoked or reflex seizure in an individual with a greater than 60% risk of experiencing another seizure within the next 10 years, or as an epilepsy syndrome (Fisher et al., 2014). The primary manifestation of epilepsy is the occurrence of epileptic seizures, which are transient events characterized by abnormal neuronal activity in the brain, leading to signs and/or symptoms. These seizures are often accompanied by stereotyped behavioral changes that reflect the underlying neural mechanisms of the disease (Beghi, 2020; Fisher et al., 2005).

The primary treatment for epilepsy involves the use of pharmacological interventions aims to reduce or prevent seizure occurrences. However, approximately 80% of patients experience adverse effects or side effects from antiseizure medications, which can significantly impact their quality of life and lead to treatment interruption or non-adherence (Devinsky et al., 2018). Moreover, despite the existence of more than 20 antiseizure drugs, around 30% of individuals with epilepsy do not achieve seizure control with these medications (Stafstrom & Carmant, 2015). Recent systematic reviews with meta-analyses have reported a cumulative incidence of drug-resistant epilepsy at 25% in studies involving children and 14.6% in studies involving adults or mixed-age populations (Sultana et al., 2021). Consequently, there is a crucial need to explore new treatment approaches and develop novel antiepileptic drugs to effectively manage epilepsy (Bialer & White, 2010).

N-acetylcysteine (NAC) has emerged as a promising adjunctive treatment for various neuropsychiatric disorders, including bipolar disorder, schizophrenia, obsessive-compulsive disorder, anxiety, depression, and drug addiction (Skvarc et al., 2017; Zheng et al., 2018). Additionally, NAC exhibits antioxidant, anxiolytic, and anti-stress effects in animal models, such as rodents and zebrafish (Mocelin et al., 2015, 2018; Santos et al., 2017). A recent study by Bilister Egilmez et al. (2023) demonstrated a dose-dependent reduction in seizure severity and prolonged onset time of the first myoclonic jerk in rats pretreated with NAC.

Another compound with potential therapeutic benefits is Acetyl-L-carnitine (ALC), a dietary supplement available in health food stores. ALC has shown positive effects in rodent models of depression exposed to unpredictable chronic stress (Nasca et al., 2013). Furthermore, ALC has been found to prevent PTZ-induced epileptic seizures in rats, decrease oxidative stress markers, increase glutathione (GSH) levels, and reduce the expression of apoptosis markers such as caspase-3 (Hussein et al., 2018).

The increase of glutamate neurotransmission is associated with epileptic crisis (Devinsky, et al. 2018). Both NAC and ALC have demonstrated neuroprotective effects by modulating glutamate levels (Baker et al., 2002; Moran et al., 2005; Nasca et al., 2013). These compounds hold promise as potential treatments for epilepsy, as they could potentially regulate the excitatory/inhibitory imbalance associated with the disease.

Zebrafish (*Danio rerio,* Hamilton 1822) is a small freshwater teleost species commonly used as a model organism to study epileptic seizure occurrence, the action of antiepileptic drugs, and alternative therapies for seizure control in epilepsy (Baraban et al., 2005; Siebel et al., 2015; Bertoncello et al., 2018). In this model organism, acute seizures can be induced by a chemoconvulsant pentylenetetrazole (PTZ), a gamma-aminobutyric acid (GABA)-A antagonist. PTZ reliably triggers characteristic and well-defined seizure behaviors in zebrafish, both in larvae and adult stages (Baraban et al., 2005; Mussulini et al., 2013). Additionally, zebrafish treated with antiepileptic drugs have shown a preventive effect against epileptic seizures when exposed to convulsive agents (Baraban et al., 2005; Berghmans et al., 2007).

Considering the challenges posed by epilepsy and the limitations of current therapies, this study aims to evaluate the effects of neuroprotective compounds (NAC and ALC) on acute epileptic seizures induced by PTZ in different stages of zebrafish development. The objective is to identify new alternatives for effective treatments with fewer adverse effects.

## MATERIALS AND METHODS

### Chemicals

N-acetylcysteine (NAC), acetyl-L-carnitine (ALC), and pentylenetetrazole (PTZ) compounds were obtained from Sigma-Aldrich® (CAS number: 616-91-1, 204259-54-1, 54-95-5, respectively). The diazepam (DZP) (CAS number: 439-14-5) was chosen as a positive control for the prevention of epileptic seizures.

### Animals

Adult short-fin wild-type zebrafish of both sexes were obtained from a local pet shop and acclimated for at least 2 weeks in an aquarium system with automated control of physical-chemical parameters and water recirculation (Altamar, Santos, SP) before the experiments were conducted. The following parameters were noted: temperature 28 ± 1 ° C, pH 7.5 ± 0.5, conductivity, 500 uS/cm, and 14/10 h (light/dark). The animals were housed in 14 L aquariums with a maximum density of 5 animals/L. The fish were fed twice a day with commercial feed (Poytara®, Brazil) and *Artemia* sp.

Zebrafish embryos were obtained from the natural mating of adult zebrafish maintained in static aquariums in our laboratory. For breeding, 2 male and 1 female adult zebrafish were randomly selected from home aquariums and placed in breeding tanks (approximately at 5 pm) with a grid specifically designed to prevent egg predation by separating the adults from the embryos. Females and males were separated overnight by a transparent barrier, which was removed after the lights went on the following morning approximately at 9 am. Then, approximately at 10 am, the adult animals were removed and the embryos were collected with a Pasteur pipette. Fertilized eggs were collected, washed with system water, and transferred to Petri dishes (30 larvae per dish). Embryos were maintained in a biochemical oxygen demand incubator (BOD) at 28 °C and monitored daily until 6 days post-fertilization (dpf). Since zebrafish larvae can absorb all the necessary nutrients through the yolk sac up to 7 dpf, it was not necessary to feed the animals during the experiment. The dead or abnormal embryos i.e., asymmetrical, showing coagulation, formation of vesicles, or damaged chorions were discarded.

Euthanasia of animals was performed after the tests by immersion in cold water (0 to 4° C) until the cessation of the movements, to ensure death by hypoxia according to the AVMA Guidelines for the Euthanasia of Animals (Leary et al., 2020). The protocol was approved by the Ethics Committee for Animal Use (CEUA) of Universidade Federal do Rio Grande do Sul (UFRGS) (CEUA/UFRGS n° 35823) and followed the guidelines of Conselho Nacional de Controle de Experimentação Animal (CONCEA) and experiments are reported in compliance with the ARRIVE guidelines 2.0 (du Sert et al., 2020).

### Treatments

For the tests conducted in adult animals, a total of 160 adult male and female zebrafish (50:50 ratio) were randomly selected from their home aquariums using a random number generator (random.org) to fulfill the experimental groups. The concentrations of N-acetylcysteine (NAC) and acetyl-L-carnitine (ALC) chosen for the experiment were based on previous studies by Marcon et al. (2019), Mocelin et al. (2018), and Pancotto et al. (2018). Three concentrations of treatment were used: 0.1, 1.0, and 10 mg/L. Diazepam (DZP) was used as a positive control at a concentration of 50 µM. The control group received only system water. The zebrafish were individually treated for 40 min in a beaker containing the respective concentrations of NAC, ALC, or DZP (400 mL solution), following the protocol described by Mussulini et al. (2013) and Fontana et al. (2019).

For experiments involving larvae, a total of 200 individuals were used. After breeding and daily monitoring of the embryos, until they reached 6 days post-fertilization (dpf), the larvae were randomized and placed individually in a 24-well plate, with each well containing 2 mL of the respective treatments: NAC and ALC at concentrations of 0.1, 1.0, and 10 mg/L, and DZP at a concentration of 18 µM, as the protocol established by Afrikanova et al. (2013). The control group received only system water. The duration of treatment was 18 hours.

### PTZ-induced seizures

After pretreatments, adult zebrafish were individually exposed to a 10 mM PTZ solution in an observation tank and their behavior was recorded on video for 20 min. The behavioral phenotype of PTZ-induced seizures was manually quantified from the recorded video analysis, using a standardized staged analysis model as shown in Table 1 (Mussulini et al., 2013). The highest epileptic seizure stage that each animal reached in 30-second intervals over the total analysis time (20 min) was recorded, and the latency to reach stage 4 and stage 5 of the seizure was also considered.

**Table 1.**
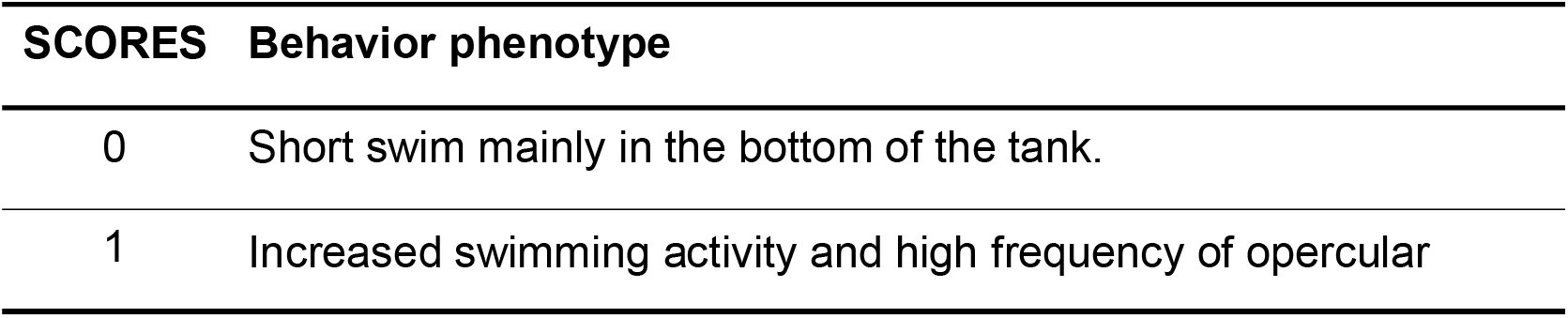
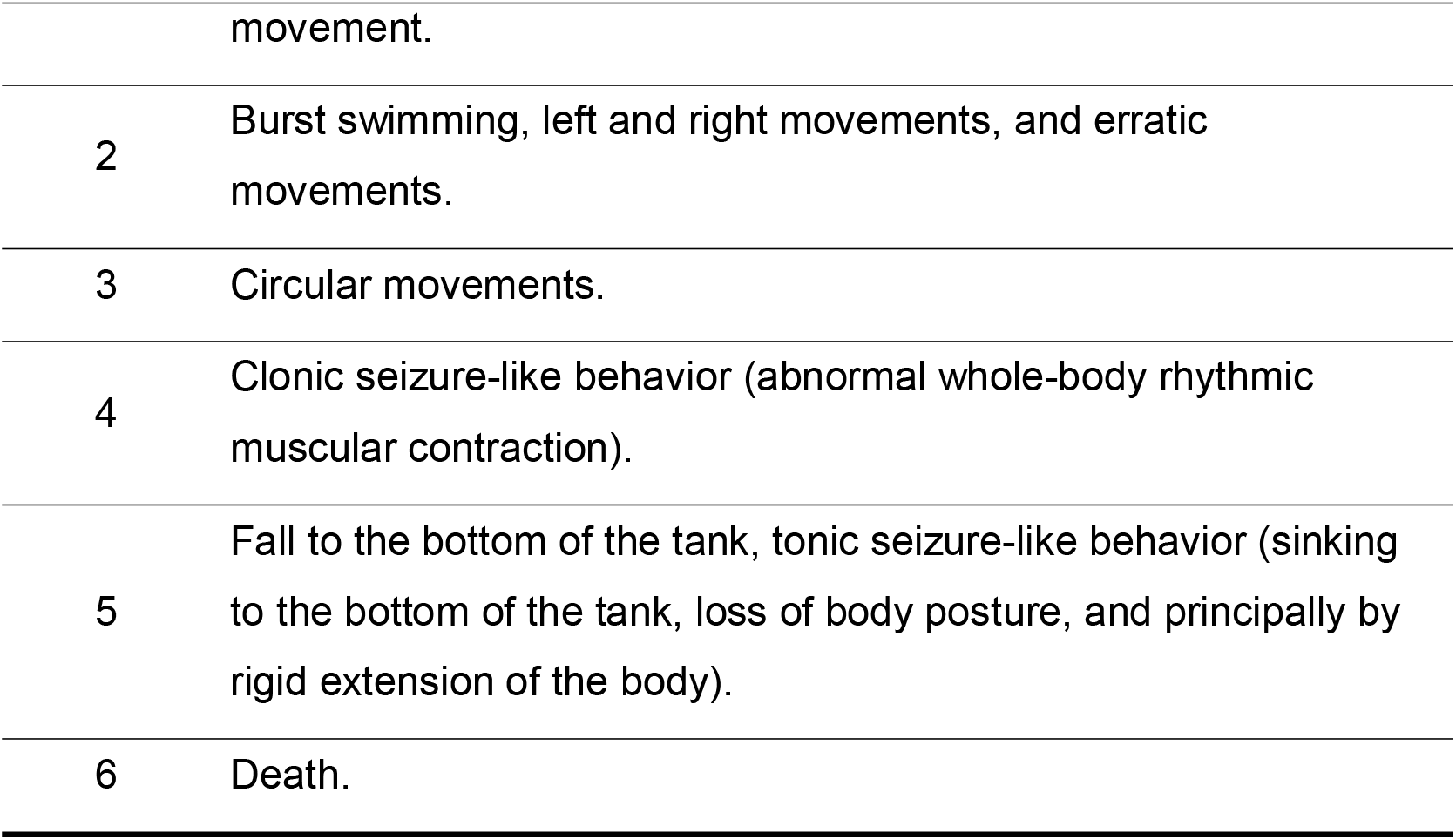
Scores and behavior phenotypes used to characterize the PTZ seizure model in adult zebrafish (Mussulini et al., 2013).

For larvae, after treatment for 18 hours, the animals (7 dpf) were individually exposed to a 10 mM PTZ solution for 10 min in a 24-well plate with 2 mL of the solution. The seizure scores were quantified using a previously standardized behavior according to Table 2 (Braban, et al. 2005) at intervals of 30 seconds, as done in adult animals. In addition, latencies for scores 2 and 3 were assessed.

**Table 2.** Scores and behavior phenotypes used to characterize the PTZ seizure model in zebrafish larvae (Baraban et al., 2005)

| **SCORES** | **Behavior phenotype** |
| --- | --- |
| 0 | Normal swim. |
| 1 | Increased swimming activity. |
| 2 | Whirlpool swimming behavior. |
| 3 | Clonus-like seizures followed by loss of posture. |

All PTZ exposures were recorded and the behavior was analyzed by researchers blinded to treatments in the software BORIS® (Friard & Gamba, 2016). Experimental groups consisted of 16 adult zebrafish and 20 larvae zebrafish. All tests were done in the morning.

### Statistical analysis

The sample size was calculated using G*Power 3.1.9.7 for Windows to detect an effect size of 0.4 with a power of 0.8 for adults and 0.9 for larvae and an alpha of 0.05 taking into account 5 experimental groups. The seizure latency was defined as the primary outcome for both groups. The total sample size was 80 for adults (n=16) and 100 for larvae (n=20) in experiments with NAC and ALC.

The normality and homogeneity of variances were confirmed for all data sets using D’Agostino-Pearson and Levene tests, respectively. Non-parametric data of seizure scores were expressed as a median. The area under the curve (AUC) and latencies to scores were represented as mean ± SD and analyzed by the one-way ANOVA followed by Dunnet’s test as post hoc. The significance level was set at p<0.05.

In the experiments with adults, one animal of the experimental group NAC 10 mg/L jumped from the tank during the analysis of seizure scores and was excluded from the results. In the experiments with NAC in larvae, two larvae from the DZP group were excluded because they died during treatment. In the experiment with ALC in larvae, three larvae from DZP and one from ALC 10 mg/L group were excluded because they were injured during manipulation.

## Results

### Effects of NAC pretreatment on zebrafish acute PTZ-induced seizures behavior

Progressive behavioral alterations were observed in PTZ adult zebrafish group, leading to the development of severe seizures with scores of 4-5, which corresponded to tonic-clonic seizures (Fig. 1A). As expected, the administration of DZP at a concentration of 50 µM resulted in a notable response, reducing seizure intensity as indicated by the area under the curve (Fig. 1B; F_4, 74_ = 8.73; p = 0.0044). Additionally, DZP pretreatment increased the latency to scores 4 and 5 onset compared to the PTZ group (Fig. 2A; F_4, 74_ = 5.475; p = 0.0013) and (Fig. 2B; F_4, 74_ = 2.725; p = 0.0298), confirming its effectiveness as a classic AED.

**Fig. 1.**
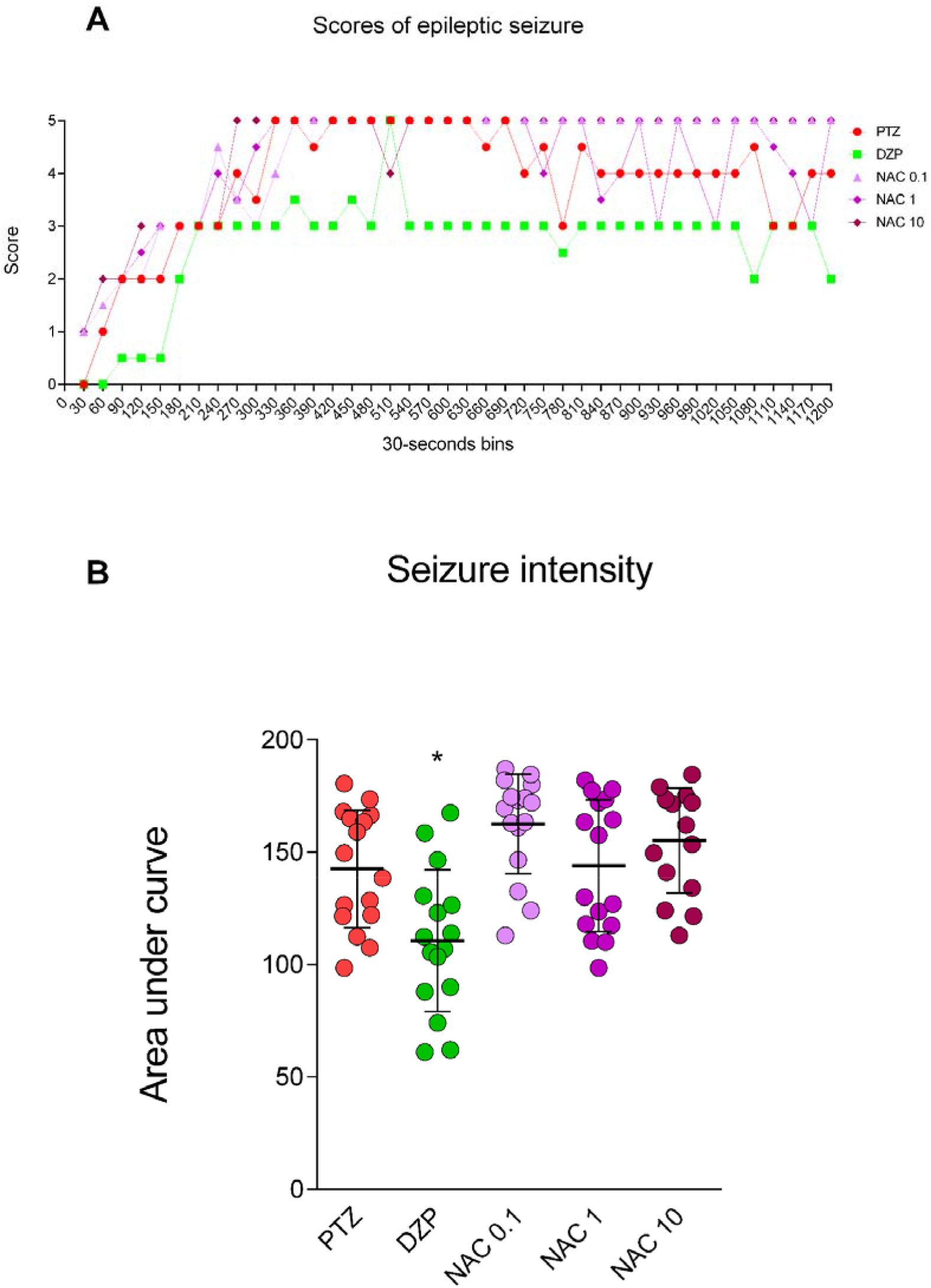
Effect of N-acetylcysteine (NAC 0.1, 1.0, and 10 mg/L) and diazepam (DZP 50 µM) on (A) scores of epileptic seizures during 20 min (only the highest score reached was consider in each interval) and (B) seizure intensity evaluated by area under the curve in adult zebrafish. Data are represented as median for scores of epileptic seizures and mean ± S.D. for seizure intensity and analyzed by one-way ANOVA followed by Dunnet’s test as post-hoc. *indicates a significant difference between groups (p<0.05). n=16, except for NAC 10 mg/L n=15

**Fig. 2.**
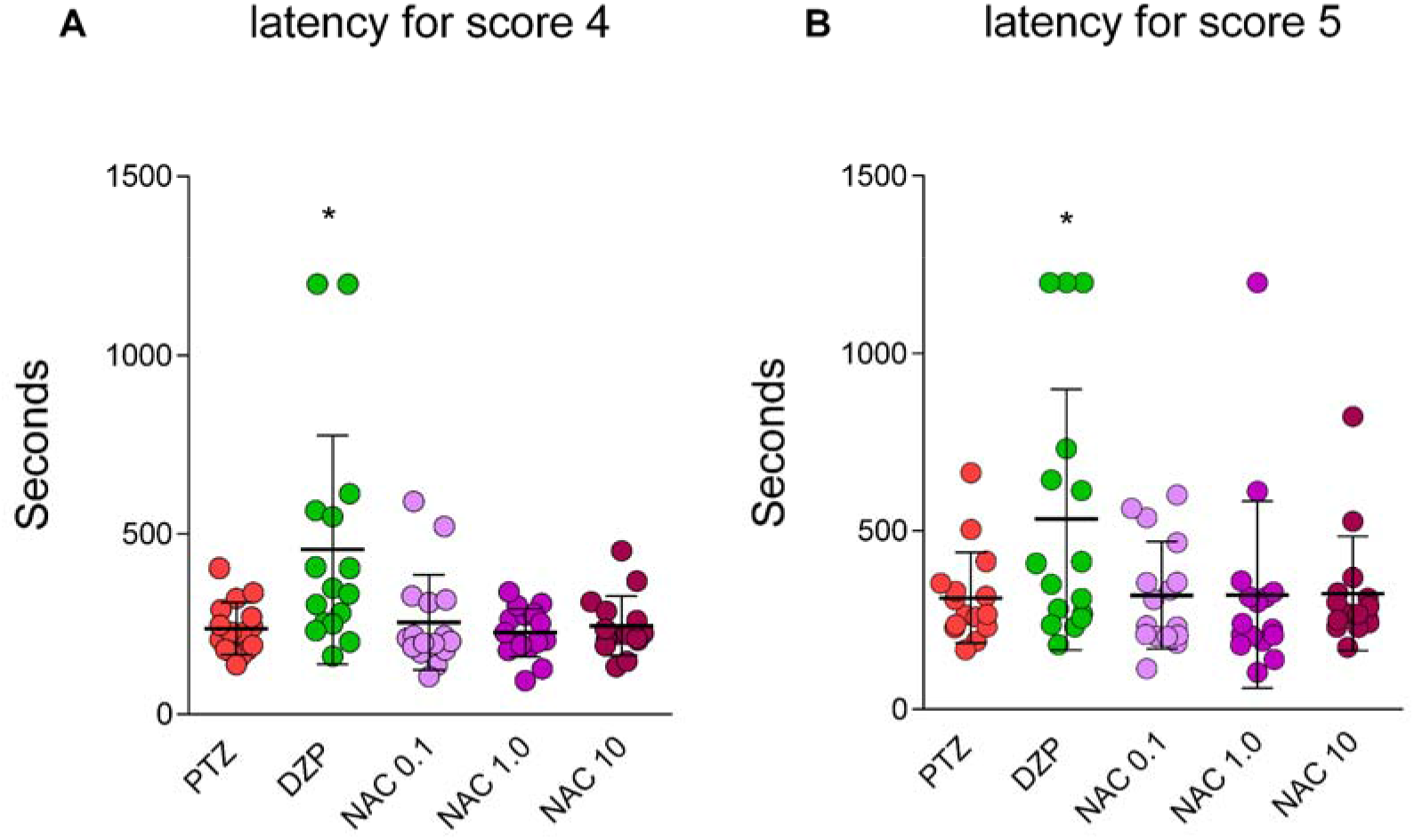
Effect of N-acetylcysteine (NAC 0.1, 1.0, and 10 mg/L) and diazepam (DZP 50 µM) in (A) latency for score 4 onset and (B) latency for score 5 onset in adult zebrafish. Data are represented as mean ± S.D. analyzed by one-way ANOVA followed by Dunnet’s test as post-hoc. *indicates a significant difference between groups (p<0.05). n=16, except for NAC 10 mg/L n=15.

On the other hand, pretreatments with NAC at concentrations of 0.1, 1.0, and 10 mg/L did not prevent or attenuate acute PTZ-induced seizures in adult zebrafish, as demonstrated by the analysis of seizure intensity (Fig. 1B) and latencies to scores 4 and 5 (Fig. 2A and B).

In larvae, we observed results similar to those observed in adults. PTZ Larvae group showed progressive seizure behavior until they reached the most severe seizure scores, a score 3 for larvae (Fig. 3 A). Pretreatment with DZP at a concentration of 18 µM effectively reduced the severity of acute PTZ-induced seizures in larvae. This was evident from the decrease in seizure intensity across the area under the curve (Fig. 3 B, F_4, 93_ = 13.68; p < 0.0001) and the increased latency observed for scores 2 and 3 of epileptic seizures in the larvae (Fig 4. A, F_4, 93_ = 11.80; p < 0.0001 and B, F_4, 93_ = 14.57; p < 0.0001). Unfortunately, despite testing three different concentrations, NAC was unable to prevent the increase in seizure scores (Fig 3 A and B) or reduce the latency for scores 2 and 3 in the larvae compared to the PTZ group (Fig. 4 A and B).

**Fig. 3.**
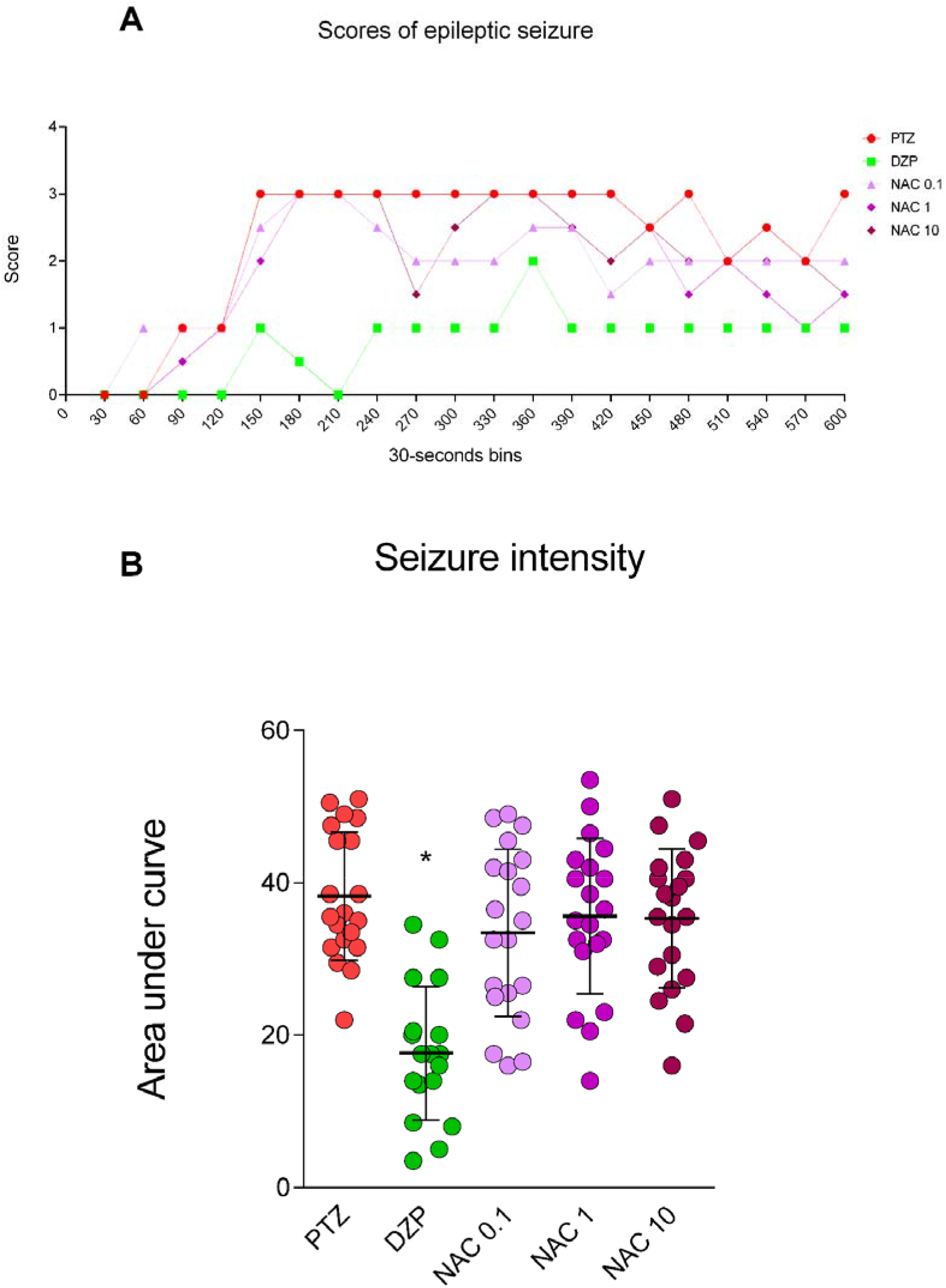
Effect of N-acetylcysteine (NAC 0.1, 1.0, and 10 mg/L) and diazepam (DZP 18 µM) in (A) Scores of epileptic seizures during 10 min (only the highest score reached was considered in each interval) and (B) seizure intensity evaluated by area under the curve in zebrafish larvae. Data are represented as a median for scores of epileptic seizures and mean ± S.D. for seizure intensity and analyzed by one-way ANOVA followed by Dunnet’s test as post-hoc. *indicates a significant difference between groups (p<0.05). n=20, except for DZP n=18.

**Fig. 4.**
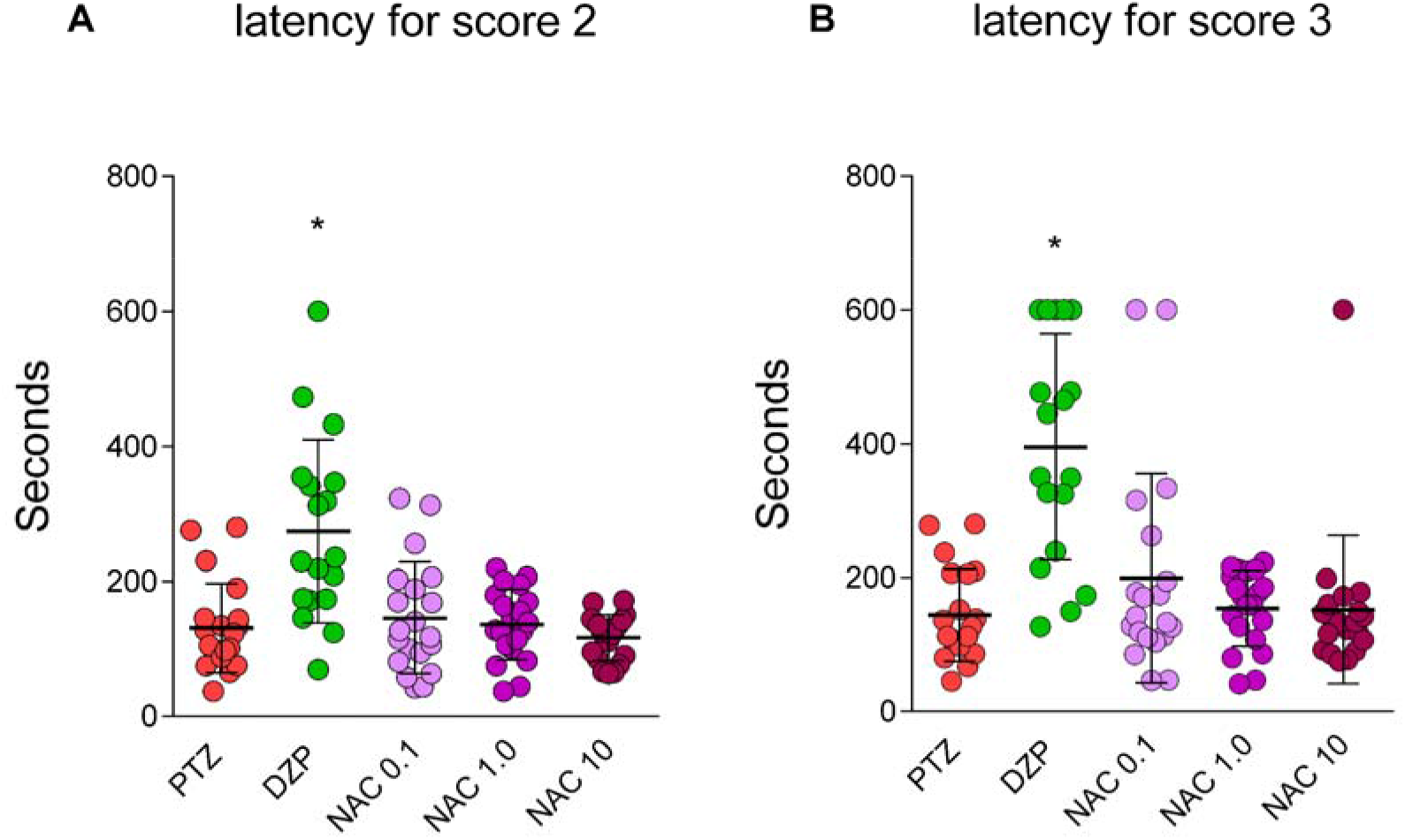
Effect of N-acetylcysteine (NAC 0.1, 1.0, and 10 mg/L) and diazepam (DZP 18 µM) in (A) latency for score 2 onset and (B) latency for score 3 onset in zebrafish larvae. Data are represented as mean ± S.D. analyzed by one-way ANOVA followed by Dunnet’s test as post-hoc. *indicates a significant difference between groups (p<0.05). n=20, except for DZP n=18

### Effects of ALC pretreatment on zebrafish acute PTZ-induced seizures behavior

In the experiments with ALC, the impact of PTZ on the animals mirrored that of the experiments with NAC. DZP demonstrated a reduction in the effect of PTZ (Fig. 5 A and B, F_4, 75_ = 6.034, p = 0.0063). However, the effects of ALC at concentrations of 0.1, 1.0, and 10 mg/L on seizures caused by PTZ were ineffective, as there was no observed decrease in seizure intensity (Fig. 5 A and B). When evaluating the latency for adult animals to reach scores 4 and 5 of epileptic seizures, ALC did not show any significant differences compared to the PTZ group at any of the tested concentrations. On the other hand, DZP at 50 µM increased the latency for scores 4 and 5 (Fig. 6 A, F_4, 75_ = 7.377, p = 0.0005 and B, F_4, 75_ = 5.119, p = 0.0132).

**Fig. 5.**
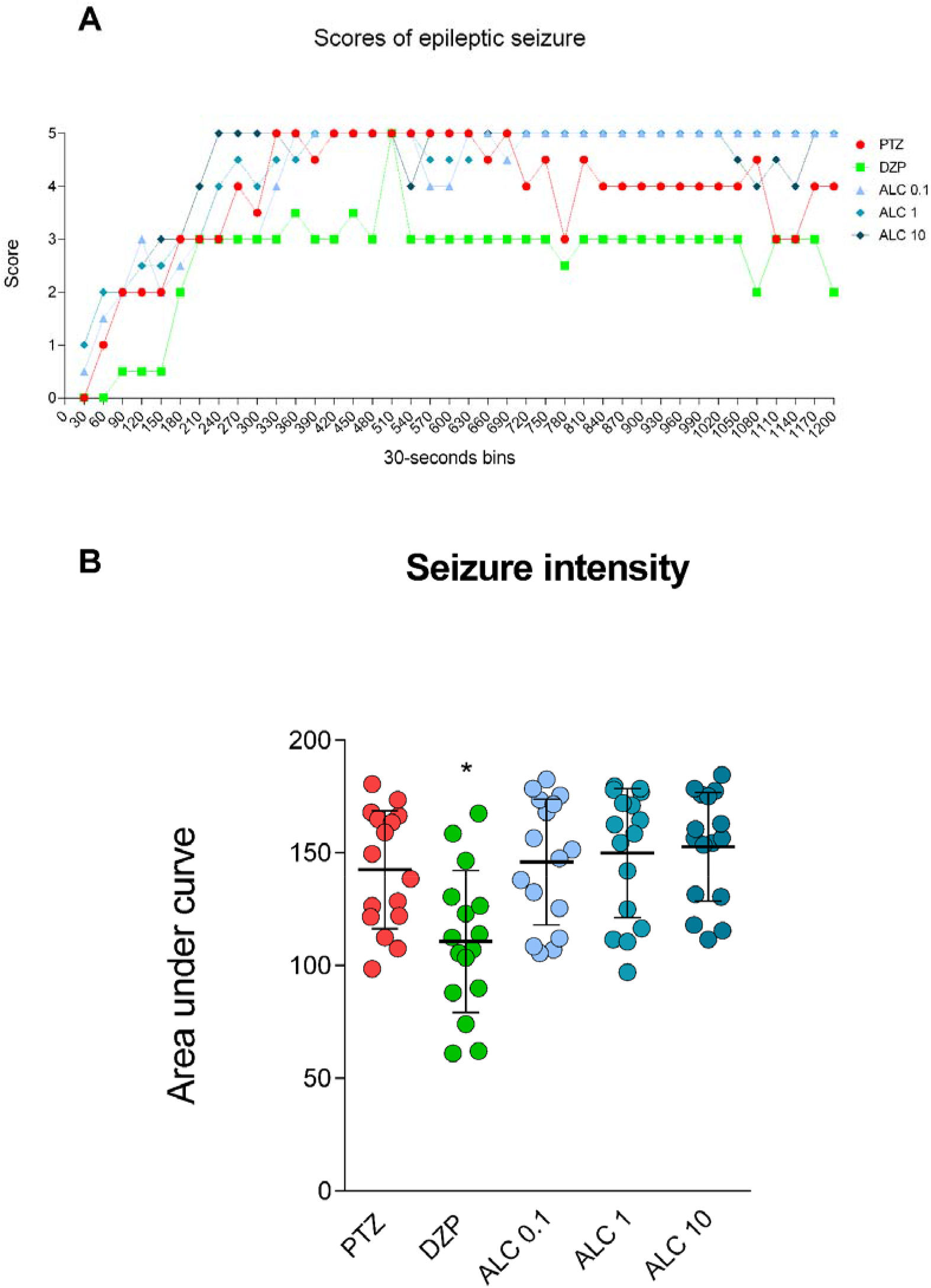
Effect of acetyl-L-carnitine (ALC 0.1, 1.0, and 10 mg/L) and diazepam (DZP 50 µM) in (A) Scores of epileptic seizures during 20 min (only the highest score reached was considered in each interval) and (B) seizure intensity evaluated by area under the curve in adult zebrafish. Data are represented as a median for scores of epileptic seizures and mean ± S.D. for seizure intensity and analyzed by one-way ANOVA followed by Dunnet’s test as post-hoc. *indicates significant difference between groups (p<0.05). n=16

**Fig. 6.**
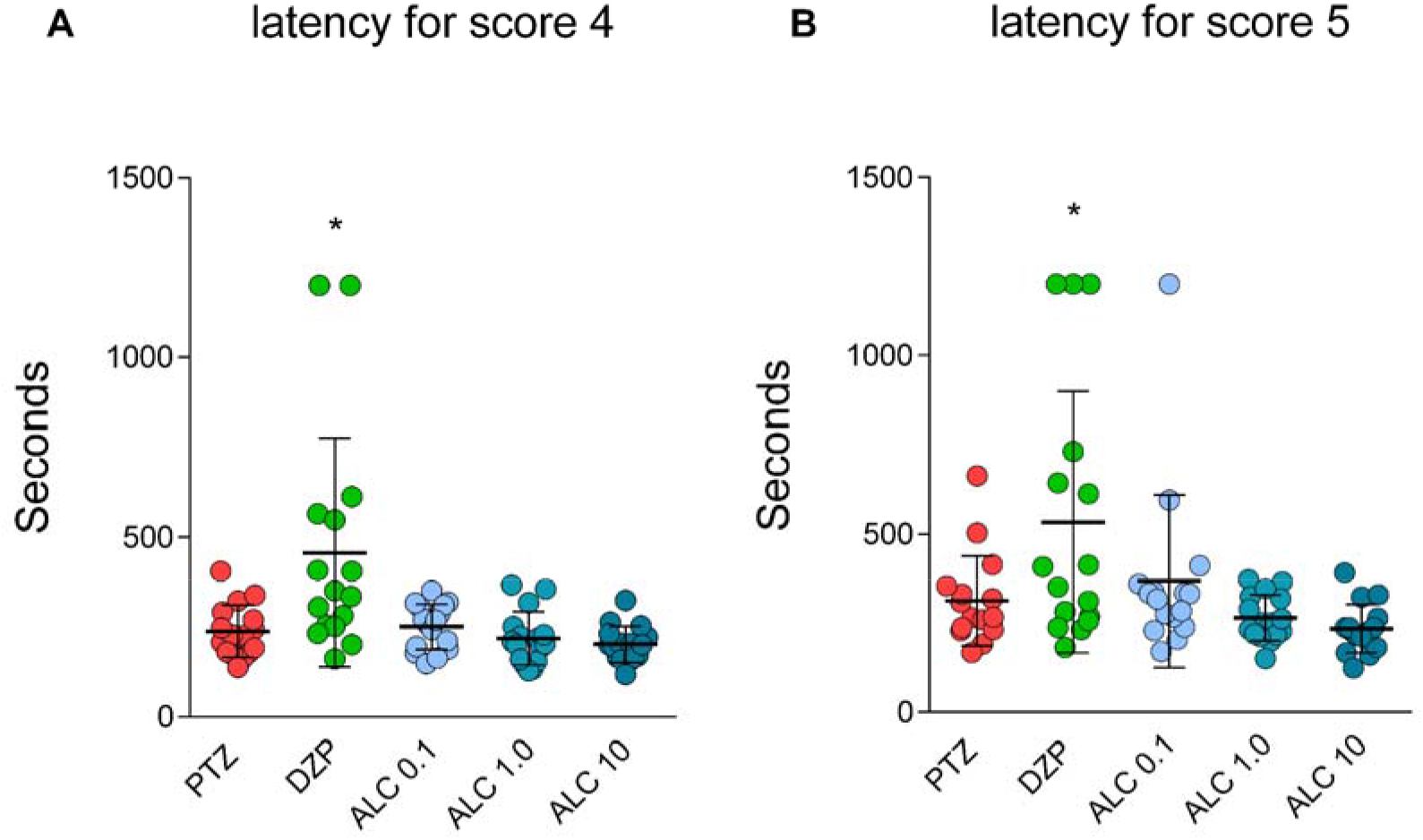
Effect of acetyl-L-carnitine (ALC 0.1, 1.0, and 10 mg/L) and diazepam (DZP 50 µM) in (A) latency for score 4 onset and (B) latency for score 5 onset in adult zebrafish. Data are represented as a mean ± S.D. analyzed by one-way ANOVA followed by Dunnet’s test as post-hoc. *indicates a significant difference between groups (p<0.05). n=16

In zebrafish larvae, the positive control diazepam at 18 µM was able to reduce the severity of the seizures (Fig. 7 A and B, F_4, 91_ = 19.38, p < 0.0001). The latencies for scores 2 and 3 increased when treated with DZP at 18 µM compared to the PTZ group. However, ALC at concentrations of 0.1, 1.0, and 10 mg/L failed to prevent the effects of PTZ on epileptic seizures, as there were no statistically significant differences in the area under the curve. Larvae treated with ALC did not show any significant differences compared to the PTZ group at any of the tested concentrations in the latencies to score 2 and 3 (Fig. 8 A, F_4, 91_ = 22.80, p < 0.0001 and B, F_4, 91_ = 16.85, p < 0.0001).

**Fig. 7.**
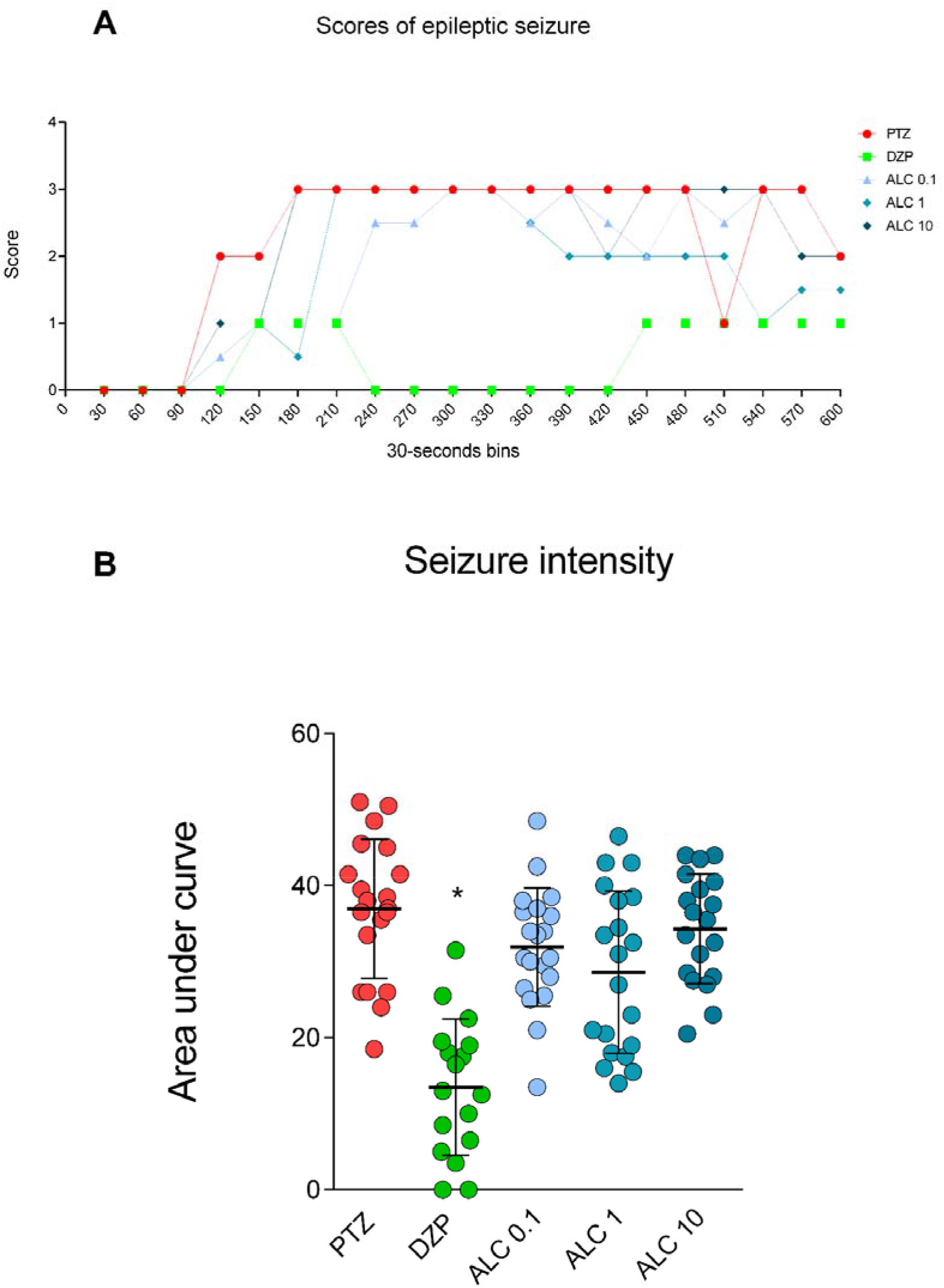
Effect of acetyl-L-carnitine (ALC 0.1, 1.0, and 10 mg/L) and diazepam (DZP 18 µM) in (A) scores of epileptic seizures during 10 min (only the highest score reached was considered in each interval) and (B) seizure intensity evaluated by area under the curve in zebrafish larvae. Data are represented as a median for scores of epileptic seizures and mean ± S.E.M. for seizure intensity and analyzed by one-way ANOVA followed by Dunnet’s test as post-hoc. *indicates a significant difference between groups (p<0.05). n=20, except for DZP group n=17, ALC 1.0 mg/L n=19 and ALC 10 mg/L n=19.

**Fig. 8.**
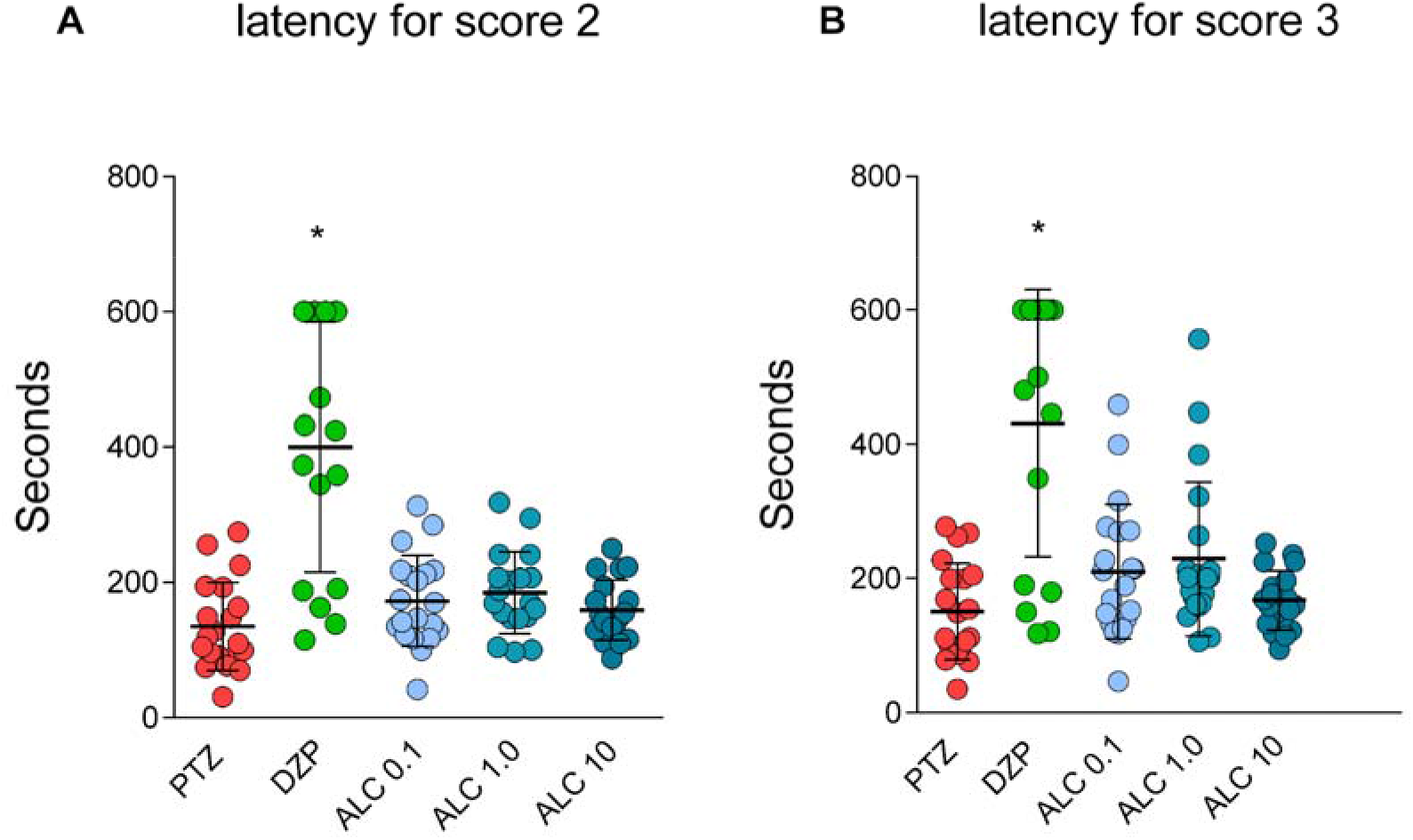
Effect of acetyl-L-carnitine (ALC 0.1, 1.0, and 10 mg/L) and diazepam (DZP 18 µM) in (A) latency for score 2 onset and (B) latency for score 3 onset in zebrafish larvae. Data are represented as mean ± S.D. analyzed by one-way ANOVA followed by Dunnet’s test as post-hoc. * indicates a significant difference between groups (p<0.05). n = 20, except DZP group n=17, ALC 1.0 mg/L n=19 and ALC 10 mg/L n=19.

## Discussion

In this study, we examined the effects of NAC and ALC on acute PTZ-induced seizures in adult and larval zebrafish. Our findings indicate that both drugs at any of the tested concentrations were not able to reduce PTZ-induced epileptic seizures in zebrafish. These molecules failed to reduce seizure intensity or increase latency for the highest seizure scores. In contrast, diazepam effectively reduced seizure intensity and increased latency for the highest scores, validating the test.

To induce acute seizures in zebrafish, we used a 10 mM concentration of PTZ. A 20-minute exposure to this concentration resulted in all seizure-like behavioral phenotypes and demonstrated lower mortality compared to a 15 mM concentration in adult zebrafish (Mussulini et al., 2013), indicating that 10 mM is a suitable concentration for studying new potential antiseizure molecules (Canzian et al., 2019; Fontana et al., 2019). Additionally, PTZ at 10 mM exhibited similar effects on larvae, accompanied by changes in electrophysiology, including increased epileptiform discharges and ictal and interictal bursts (Y. Kim et al., 2009; Y.-H. Kim et al., 2010; Hong et al., 2016). Our study using 10 mM PTZ corroborated the findings of Baraban et al. (2005), showing an increase in seizure scores over time in larvae zebrafish.

In epileptogenesis, it is known that activated astrocytes undergo molecular alterations that can contribute to neuronal hyperexcitability, such as the downregulation of gap junction connexins, glutamate transporters, and potassium channels (Devinsky et al., 2018). Impaired astrocyte function can result in decreased elimination of extracellular glutamate, leading to a reduced seizure threshold and an imbalance between neuronal excitation and inhibition (Bialer & White, 2010). Drugs that modulate glutamate, such as topiramate, perampanel, levetiracetam, and felbamate, have been used for the treatment of epileptic seizures (Sills & Rogawski, 2020). Some of these molecules have shown effects in increasing latency to seizures induced by acute PTZ administration (Pieróg et al., 2021). NAC and ALC have been reported to have glutamate-modulating effects. NAC activates cystine-glutamate antiport transporters present in astrocytes, leading to the release of glutamate that stimulates extra-synaptic metabotropic receptors (mGluRs), thereby reducing the synaptic release of the neurotransmitter (Baker et al., 2002; Moran et al., 2005). On the other hand, ALC has demonstrated effects in models of depression in rodents exposed to unpredictable chronic stress through a possible epigenetic regulation of metabotropic glutamate type 2 (mGlu2) receptors. This was observed after 3 days of treatment (Nasca et al., 2013).

Several studies have demonstrated the antiseizure effects of NAC in rodents. Uma Devi et al. (2006) studied the effect of NAC (50 and 100 mg/kg for 8 days) on PTZ-induced seizures in mice (60 mg/kg via intraperitoneal injection). NAC significantly prolonged the latencies to jerks and clonic generalized seizures during the 30-minute analysis, with this effect not being dose-dependent. Associating NAC with sodium valproate resulted in a significant enhancement of the anticonvulsant effect. Zaeri & Emamghoreishi (2015) also demonstrated the antiseizure effect of NAC (50-150 mg/kg) in an acute and chronic PTZ seizure model (90 mg/kg via intraperitoneal injection). In our study, we did not observe an antiseizure effect of NAC at concentrations of 0.1, 1.0, and 10 mg/L when administered for 40 minutes (adults) or 18 hours (larvae) in the PTZ seizure model in zebrafish. This lack of effect may be attributed to the low concentrations used in our study and the duration of treatment (acute vs. chronic in other studies). It is important to note that a study by Tallarico et al. (2022) demonstrated that NAC treatment significantly increased the number and duration of spike-wave discharges in WAG/Rij rats (rat model of absence epilepsy), exacerbating absence epilepsy while ameliorating neuropsychiatric comorbidities. Therefore, the effect of NAC on seizures may depend on the seizure type.

ALC also did not demonstrate an antiseizure effect on acute PTZ-induced seizures in zebrafish, despite other researchers have demonstrated the antiseizure effect in rodents using the kindling model. The kindling model involves repeated electrical stimuli of sub-effective intensity or repeated administration of a low dose of a chemoconvulsant until complete tonic-clonic seizures are induced (Mason & Cooper, 1972; Dhir, 2012; Davoudi et al., 2013). Hussein et al. (2018) exposed animals to PTZ (40 mg/kg i.p., 3 times/week, for 3 weeks) and divided them into two groups. The first group received oral L-carnitine (100 mg/kg/day for 4 weeks), while the second group received saline. L-carnitine treatment decreased seizure scores and duration, increased latency to the first seizure, attenuated PTZ-induced increase in the level of the oxidative stress marker malondialdehyde (MDA), and increased antioxidant glutathione (GSH) activity. More recently, Essawy et al. (2022) also demonstrated the antiseizure effect of L-carnitine (300 mg/kg) in a rodent kindling model. Pre- and post-treatment with L-carnitine suppressed the kindling acquisition process and remarkably alleviated the PTZ-induced effects. Additionally, a study showed the antiseizure effect of ALC (100 mg/kg) in the kainate murine model of temporal lobe epilepsy by attenuating status epilepticus and reducing oxidative stress and neuroinflammation (Tashakori-Miyanroudi et al., 2022).

Diazepam, a benzodiazepine medication, is commonly used to treat acute recurrent seizures by enhancing the activity of gamma-aminobutyric acid (GABA) (Dhaliwal et al., 2023). Our study showed that diazepam effectively increased latency for the highest seizure scores and reduced seizure intensity at concentrations of 50 µM (adults) and 18 µM (larvae) in zebrafish. These results support other studies where diazepam was used as a positive control to reduce seizures induced by PTZ in zebrafish (Choo et al., 2018; da Silva et al., 2021).

As prospects, further studies can be conducted to evaluate the effect of acute treatment with NAC and ALC in higher concentrations on PTZ-induced seizures. Furthermore, NAC and ALC could be tested as adjuncts to other antiseizure treatments to determine if they enhance the overall treatment effect. Additionally, the kindling model is not well standardized in zebrafish, as the existing studies only exposed the animals to chronic low doses of PTZ and evaluated the seizure behavior just after the exposure. Spontaneous seizures do not occur in existing protocols (Kumari et al., 2020; Kundap et al., 2017), therefore NAC and ALC could be evaluated in a new kindling model in zebrafish to assess their antiseizure effects.

## Conclusion

In this study, we aimed to investigate the potential antiseizure effects of NAC and ALC through the utilization of an acute PTZ-induced seizure model in adult and larvae zebrafish. Our findings indicate that acute treatment with both drugs did not exhibit any significant antiseizure effects. The external validity of previous findings from rodent experiments is thus limited. Nevertheless, it is important to note that further investigations are warranted to fully assess the effects of NAC and ALC on PTZ-induced epileptic seizures. Potential approaches for future research include the exploration of higher concentrations of NAC and ALC, extended treatment durations, and the incorporation of electrophysiological evaluation into the experimental design. By conducting these additional studies, it will be possible to more comprehensively understand the potential antiseizure properties of NAC and ALC in the context of PTZ-induced seizures.

## AUTHOR CONTRIBUTION

Conceptualization: RC and AP; Methodology: RC, CGR, TSB, AL, RB, MM, AP; Investigation: RC, CGR, TSB, AL, RB, MM; Writtin-Original Draft: RC; Writting-Review and Editing: RC, CGR, TSB, AL, RB, MM, APH, AP; Supervision: AP; Funding Acquisition: AP.

## ACKNOWLEDGMENTS

We thank the Conselho Nacional de Desenvolvimento Científico e Tecnológico (CNPq, proc. 303343/2020-6), Coordenação de Aperfeiçoamento de Pessoal de Nível Superior - Brasil (CAPES), and Pró-Reitoria de Pesquisa (PROPESQ) at Universidade Federal do Rio Grande do Sul (UFRGS) for funding and support.

